# Distinct and Complementary Functions of Rho kinase isoforms ROCK1 and ROCK2 in Prefrontal Cortex Structural Plasticity

**DOI:** 10.1101/275404

**Authors:** Kelsey M. Greathouse, Benjamin D. Boros, Josue F. Deslauriers, Benjamin W. Henderson, Kendall A. Curtis, Erik G. Gentry, Jeremy H. Herskowitz

**Affiliations:** Center for Neurodegeneration and Experimental Therapeutics and University of Alabama at Birmingham, Birmingham, Alabama 35294; Department of Neurology, University of Alabama at Birmingham, Birmingham, Alabama 35294

## Abstract

Twenty-nine protein kinase inhibitors have been used to treat human diseases. Out of these, two are Rho-associated protein kinase (ROCK) 1 and 2 inhibitors. ROCKs are attractive drug targets for a range of neurologic disorders; however a critical barrier to ROCK-based therapeutics is ambiguity over whether there are isoform-specific roles for ROCKs in neuronal structural plasticity. Here, we used a genetics approach to address this long-standing question. Both male and female adult ROCK1^+/−^ and ROCK2^+/−^ mice exhibited anxiety-like behaviors compared to littermate controls. Individual pyramidal neurons in the medial prefrontal cortex (mPFC) were targeted for iontophoretic microinjection of fluorescent dye, followed by high-resolution confocal microscopy and neuronal 3D reconstructions for morphometry analysis. Increased apical and basolateral dendritic length and intersections were observed in ROCK1^+/−^ but not ROCK2^+/−^ mice. Although dendritic spine densities were comparable among genotypes, apical spine extent was decreased in ROCK1^+/−^ but increased in ROCK2^+/−^ mice. Spine head and neck diameter were reduced similarly in ROCK1^+/−^ and ROCK2^+/−^ mice; however certain spine morphologic subclasses were more affected than others in a genotype-dependent manner. Biochemical analyses of ROCK substrates revealed that phosphorylation of LIM kinase was reduced in synaptic fractions from ROCK1^+/−^ or ROCK2^+/−^ mice, correlating to overlapping spine morphology phenotypes. Collectively, these observations implicate ROCK1 as a novel regulatory factor of neuronal dendritic structure and detail distinct and complementary roles of ROCKs in mPFC dendritic spine structural plasticity. This study provides a fundamental basis for current and future development of isoform-selective ROCK inhibitors to treat neurologic disorders.

**Significance Statement:** The Rho-associated protein kinases (ROCK) 1 and 2 heavily influence neuronal architecture and synaptic plasticity. ROCKs are exciting drug targets and pan-ROCK inhibitors are clinically approved to treat hypertension, heart failure, glaucoma, spinal cord injury, and stroke. However development of isoform-specific ROCK inhibitors is hampered due to ambiguity over ROCK1- or ROCK2-specific functions in the brain. Our study begins to address this critical barrier and demonstrates that ROCK1 can mediate the dendritic arbor of neurons while both ROCK1 and ROCK2 heavily influence dendritic spine morphology. This study highlights distinct and complementary roles for ROCK1 and ROCK1 in prefrontal cortex structural plasticity and provides a fundamental basis for future development of isoform-selective ROCK inhibitors to treat neurologic disorders.

## Introduction

Originally isolated as GTP-bound RhoA interacting proteins, the Rho-associated coiled-coil containing kinases (ROCK) are members of the AGC family of serine/threonine kinases and are extensively studied regulators of actin–myosin-mediated cytoskeleton contractility (Leung et al., 1995; Ishizaki et al., 1996; Leung et al., 1996; Matsui et al., 1996; Nakagawa et al., 1996). Two mammalian ROCK isoforms exist, ROCK1 and ROCK2, and share 65% similarity in their amino acid sequences and 92% identity in their kinase domains (Nakagawa et al., 1996). ROCK1 and ROCK2 expression patterns are largely similar in humans with higher transcript levels of ROCK1 in thymus and blood and ROCK2 in brain (Julian and Olson, 2014). ROCKs phosphorylate a number of substrates predominantly tied to cellular morphology, adhesion, and motility. These actions implicate ROCK1 and ROCK2 as putative therapeutic targets for a variety of human conditions, such as cancer, asthma, insulin resistance, kidney failure, osteoporosis, and erectile dysfunction (Olson, 2008; Schaafsma et al., 2008; Lee et al., 2009; Albersen et al., 2010; Komers et al., 2011; Rath and Olson, 2012). Moreover, studies have identified pathogenic roles for ROCKs or explored the potential to repurpose ROCK inhibitors in neurologic disorders, including glaucoma, spinal cord injury, stroke, Alzheimer’s disease, Frontotemporal Dementia, Parkinson’s disease, and Amyotrophic Lateral Sclerosis (Shibuya et al., 2005; Duffy et al., 2009; Herskowitz et al., 2013; Challa and Arnold, 2014; Koch et al., 2014; Gentry et al., 2016; Henderson et al., 2016; Tatenhorst et al., 2016; Gunther et al., 2017).

Pharmacologic studies have driven much of our understanding of ROCKs in the brain, with Fasudil and Y-27632 being the most widely characterized inhibitors. However these and other current ROCK inhibitors are not isoform-specific and likely inhibit other kinases, including PKA and PKC, at higher doses used for *in vivo* experiments (Davies et al., 2000). Therefore, it is challenging to assign functions to ROCK1 or ROCK2 based on the kinase inhibitors. Despite these caveats, ROCK-based drug studies continue to extend the implications of this treatment strategy for brain disorders as well as illuminate basic functions of the ROCKs. For instance, recent findings indicate that directed delivery of Fasudil to the prefrontal cortex enhances goal-directed behavior in mice and blocks habitual response for cocaine (DePoy et al., 2017; Swanson et al., 2017). Pharmacologic inhibition of ROCKs as a therapeutic for adolescent or adult drug addiction is an exciting hypothesis, yet these applications continue to raises important questions about the contribution of ROCK1 or ROCK2 to the observed beneficial effects of pan-ROCK inhibitors.

Although ROCKs share protein substrates, including myosin light chain (MLC), myosin light chain phosphatase, and LIM kinases (LIMK), evidence from genetic approaches in cell-based assays suggests distinct functions of ROCK isoforms (Amano et al., 1996; Kimura et al., 1996; Sumi et al., 2001). Older studies from homozygous knockout mice revealed major developmental problems or embryonic lethality in ROCK1^-/−^ or ROCK2^-/−^, respectively; however, a different genetic background alleviated some of the effects from ROCK1 deletion (Thumkeo et al., 2003; Shimizu et al., 2005; Zhang et al., 2006). Working with ROCK1^-/−^ or ROCK2^-/−^ mouse embryonic fibroblasts, Shi et al. studied differential roles for ROCK1 and ROCK2 in regulating actin cytoskeleton reorganization after doxorubicin exposure. These findings suggested ROCK1 destabilizes actin via MLC phosphorylation whereas ROCK2 stabilizes actin through cofilin phosphorylation (Shi et al., 2013).

Genetic exploration of ROCK1 and ROCK2 function in brain has been limited due to the complications of homozygous knockout mice on mixed backgrounds. Yet despite reports indicating that ROCK1^+/−^ and ROCK2^+/−^ mice develop normally, *in vivo* studies of the heterozygous models are rare (Zhang et al., 2006; Duffy et al., 2009). To this end, we independently generated new ROCK1^+/−^ and ROCK2^+/−^ mice on the C57BL/6N background to define ROCK isoform-specific functions related to structural plasticity. The studies herein provide novel distinct yet complimentary roles for ROCK1 and ROCK2 in the prefrontal cortex with implications at the molecular, cellular, and organismal level.

## Materials and Methods

### Animals

All experimental procedures were performed under a protocol approved by the Institutional Animal Care and Use Committee at the University of Alabama at Birmingham (UAB). Generation of ROCK1^+/−^ mice are described as follows: C57BL/6N-Rock1<tm1b(NCOM)Mfgc>/Tcp were made as part of the NorCOMM2 project with C57BL/6N-Rock1<tm1a(NCOM)Mfgc>/Tcp mice made from NorCOMM embryonic stem (ES) cells at the Toronto Centre for Phenogenomics (Bradley et al., 2012). C57BL/6N-Rock1<tm1b(NCOM)Mfgc>/Tcp mice were obtained from the Canadian Mouse Mutant Repository. ROCK2^+/−^ mice were generated as follows: C57BL/6N-Rock2tm1a(KOMP)Wtsi mice were made from ES cells purchased from the International Mouse Phenotyping Consortium at the University of California, Davis. ES cell injections were performed by the UAB Transgenic & Genetically Engineered Models Core. The C57BL/6N-Rock2tm1a(KOMP)Wtsi ES cells used for this research project were generated by the trans-NIH Knock-Out Mouse Project (KOMP) and obtained from the KOMP Repository (www.komp.org). NIH grants to Velocigene at Regeneron Inc (U01HG004085) and the CSD Consortium (U01HG004080) funded the generation of gene-targeted ES cells for 8500 genes in the KOMP Program and archived and distributed by the KOMP Repository at UC Davis and CHORI (U42RR024244). For more information or to obtain KOMP products go to www.komp.org or.

### Behavior

All mice were kept in a facility with a 12 hour light cycle and a 12 hour dark cycle. All behavioral testing was performed during the light cycle. Mice were placed in the testing room at a minimum of one hour before testing for acclimation. Each apparatus was disinfected with 2% chlorhexidine prior to testing. Each apparatus was cleaned with 70% after each experiment. All testing was conducted at the same time each day on consecutive days.

#### Elevated Plus Maze

The elevated plus maze apparatus (EPM; Med Associates) was 1 m high with 2 in wide arms. Two opposite arms had 8 in high black walls, while the other opposing arms were open. Each mouse was placed in the center of the maze and freely explored for 5 minutes. Exploration into arms was recorded and traced by the manufacture’s software (CleverSys). Percent time in open arms was calculated by dividing the time in open arms by total time.

#### Open Field Test

For open field assessment, mice were placed into a 16 in x 16 in plexiglass box (Med Associates) with opaque walls. Mice explored for 10 minutes, and ambulatory distance and ambulatory counts were determined by the manufacture’s software (CleverSys).

#### Y-maze

Y-maze testing was conducted as previously described (Li et al., 2014). The Y-maze consisted of three arms (38.1 cm long, 8.9 cm wide, 12.7 cm high) made of white plexiglass with randomly placed visual cues in each arm. Mice were placed in the center of the maze and allowed to explore for 5 minutes. Activity was recorded and tracked with video tracking software (Cleversys). An alternation was defined as sequential entries into each arm without re-entry into the previously explored arm. The percent of correct alternations was calculated by dividing the total number of spontaneous alternations by the total number of arm entries.

### Perfusions and tissue processing

#### PFA, vibratome, storage

Animals were anesthetized with Fatal Plus (Vortech Pharmaceuticals, Catalog #0298-9373-68). The abdominal cavity and pericardium were carefully opened to expose the heart and mice were transcardially perfused with cold 1% paraformaldehyde (PFA) for 1 min, followed by 10 min of 4% paraformaldehyde (Sigma Aldrich, Catalog #P6148) with 0.125% glutaraldehyde (Fisher Scientific, Catalog #BP2547). Peristaltic pump (Cole Parmer) was used for consistent administration of PFA. Immediately following perfusion, the mouse was decapitated and the whole brain was removed and drop fixed in 4% PFA containing glutaraldehyde for 8-12 hours at 4°C. After post-fixation, the brains were sliced in the order they were perfused coronally in 250 μm increments using a Leica vibratome (VT1000S) with a speed of 70, and frequency of 7. The platform was filled with cold 0.1 M PB buffer and the brain was glued (Loctite) perpendicular to the stage, cerebellum side down. All slices were stored one slice per well in a 48-well plate containing 0.1% sodium azide (Fisher, Catalog#BP922I) in 0.1M PB. Plates covered in parafilm are stored long term at 4°C. Notably, these procedures were performed according to (Dumitriu et al., 2011).

#### PBS, storage

Animals were anesthetized with Fatal Plus. The abdominal cavity and pericardium were carefully opened to expose the heart. Mice were transcardially perfused with cold 1X PBS for 2 minutes. Immediately following perfusion, the brain was extracted and dissected into two hemispheres. Each hemisphere was immediately flash frozen in 2-methylbutane (Sigma, Catalog#320404), placed on dry ice, and transferred to the -80°C for storage.

### Iontophoretic microinjection of fluorescent dye

Microinjections were executed using previously described methods (Dumitriu et al., 2011). We used a Nikon Eclipse FN1 upright microscope with a 10X objective and a 40X water objective placed on an air table. We used a tissue chamber assembled in the lab consisting of a 50x75 mm plastic base with a 60x10 mm petri dish epoxied to the base. A platinum wire is attached in a way that the ground wire can be connected to the bath by an alligator clip. The negative terminal of the electric current source was connected to the glass micropipette filled with 2 μl of 8% Lucifer yellow dye (ThermoFisher, Catalog#L453). Micropipettes (A-M Systems, Catalog #603500) with highly tapered tips were pulled fresh the day of use. The manual micromanipulator was secured on the air table with magnets that provided a 45° angle for injection. Using a brush, the brain slice was placed into a small petri dish containing 1X PBS and 4’,6-diamidino-2-phenylindole (DAPI) for 5 min at room temperature. After incubation in DAPI, the slice was placed on dental wax and then a piece of filter paper was used to adhere the tissue. The filter paper was then transferred to the tissue chamber filled with 1X PBS and weighted down for stability. We used the 10X objective to advance the tip of the micropipette in XY and Z until the tip is just a few micrometers above the tissue. The 40X objective was then used to advance to tip into layers II and III of the prefrontal cortex. Once the microelectrode contacted a neuron, 2 nA of negative current was used for 5 min to fill the neuron with lucifer yellow. After the 5 min, the current was turned off and the micropipette was removed from the neuron. Neuron impalement within layers II and III occurs randomly in a blind manner. If the entire neuron does not fill with dye after penetration, the electrode is removed and the neuron is not used for analysis. Multiple neurons were injected in each hemisphere of the medial prefrontal cortex of each animal. After injection the filter paper containing the tissue was moved back into the chamber containing 1X PBS. Using a brush, the tissue was carefully lifted off the paper and placed on a glass slide with two 125 μm spacers (Electron Microscopy Sciences, Catalog #70327-20S). Excess PBS was carefully removed with a Kimwipe, and the tissue was air-dried for 1 min. One drop of Vectashield (Vector Labs, Catalog #H1000) was added directly to the slice, the coverslip (Warner, Catalog #64-0716) was added, and was sealed with nail polish. Injected tissue was stored in 4°C.

### Confocal Microscopy

Confocal microscopy was used to capture images of pyramidal cells and dendrites from prefrontal cortex layers 2 and 3 and our methods were based on (Dumitriu et al., 2011). All imaging was performed by a blinded experimenter. Images were captured with a Leica LAS AF TCS MP5, Leica Microsystems 2.6.3, and either 20X or 63X oil immersion objective (Leica HCX PL Apo CS, N.A. 1.40). The following parameters were used for all images: Argon laser: 20% power; 458 Laser: 100% power; smart offset: 1.5%; smart gain: 800 V; Pinhole: 1 airy unit; image size: 1024 × 1024px. The experimenter focused on each dye-impregnated neuron and captured three-dimensional z-stacks of those meeting criteria. Criteria for imaging of neuronal arbors included: (1) located within 80 μm working distance of microscope, (2) fully impregnated with dye, (3) unobstructed by any artifact. For each neuron, z-stacks were captured with the following parameters: z-step: 0.503 μm; image size: 1024 × 1024 px (0.223 μm × 0.223 μm × 0.503 μm); zoom: 3x; line averaging: 2; acquisition rate: 700 Hz. Image stacks from were collected in.lif format. The experimenter identified secondary dendrites from dye-impregnated neurons and captured three-dimensional z-stacks of those meeting criteria. Criteria for imaging neuronal dendrites included: (1) within 80 μm working distance of microscope; (2) relatively parallel with the surface of the coronal section; (3) no overlap with other branches. Secondary dendrites of pyramidal arbors were viewed and selected at low magnification. This segment was then viewed and imaged at 63X magnification. For each neuronal dendrite, z-stacks were captured with the following parameters: z-step: 0.1259 μm; image size: 1024 × 1024 px (0.0501 μm × 0.0501 μm × 0.0501 μm); zoom: 4.8x; line averaging: 4; acquisition rate: 400 Hz. Images were registered in ImageJ using linear stack alignment with Scale Invariant Feature Transform (SIFT) and recommended settings. Registered images were saved in .tif format. Captured images were deconvolved using Huygens Deconvolution System (16.05, Scientific Volume Imaging, the Netherlands) and the following settings: GMLE; maximum iterations: 10; signal to noise ratio: 15; quality: 0.003. Prior to deconvolution, imaging meta-data from original .lif files were extracted to Huygens template files in .hgsm format and attributed to each corresponding registered image. Deconvolved images were saved in .tif format.

### Neuronal reconstructions

Neuronal arbor and dendritic spine reconstruction and analysis were performed with Neurolucida 360 (2.70.1, MBF Biosciences, Williston, Vermont). Dendritic spine analysis was performed as previously described with the noted adjustments (Boros et al., 2017). Briefly, image stacks of neuronal dendrites were imported to Neurolucida 360, and the full dendrite length was traced with semi-automatic directional kernel algorithm. The experimenter manually confirmed that all assigned points matched dendrite diameter and position in x, y, and z planes and adjusted each reconstruction if necessary. Dendritic spine reconstruction was performed automatically using a voxel-clustering algorithm and the following parameters: outer range: 5.0 μm; minimum height: 0.3 μm; detector sensitivity: 80%; minimum count: 8 voxels; backbone length. Next, the experimenter manually verified that the classifier correctly identified all protrusions. When necessary, the experimenter added any semi-automatically by increasing detector sensitivity. The morphology and backbone points of each spine were verified to ensure a representative spine shape, and merge and slice tools corrected inconsistencies. Each dendritic protrusion was automatically classified as a dendritic filopodium, thin spine, stubby spine, or mushroom spine based on morphological measurements using previously described (Boros et al., 2017). Reconstructions were collected in Neurolucida Explorer (2.70.1, MBF Biosciences, Williston, Vermont) for branched structure analysis, and then exported to Microsoft Excel (Redmond, WA). Spine density was calculated as the number of spines per μm of dendrite length. To characterize neuronal arbors, image stacks were imported to Neurolucida 360. First, the soma was detected in three-dimensions semi-automatically using sensitivity at 40-50%. Neuronal arbors were automatically detected using the following parameters: seeding: medium density; smallest segment: 30 μm; sensitivity: 75-100%. Then, the experimenter closely examined the accuracy of each arbor reconstruction in each x, y, and z dimension. When necessary, the experimenter semi-automatically joined, separated, or extended segments using detach, connect, and remove tools. The experimenter ensured that the base of each arbor was as near to the soma as possible. All extensions from the soma were manually classified as apical dendrites, basal dendrites, or axons. Apical and basal arbor reconstructions were imported to Neurolucida Explorer and separately analyzed. Sholl analysis characterized the number of intersections or length using 10 μm concentric shells (radii), and these results were exported and collected in Microsoft Excel. The number of intersections and length between Sholl radii were calculated.

### Biochemistry

#### Brain region subdissection

Hemibrains were bathed in a petri dish of ice-cold PBS with protease (Sigma, Catalog #S8820) and phosphatase inhibitors (Thermo Scientific, Catalog #1861277). Using a dissecting microscope, brain anatomical atlas, and surgical instruments, the medial prefrontal cortex was isolated from each hemibrain. All brain regions were stored at -80°C.

#### Synaptosomal preparations

Synaptosomes were prepared using the following biochemical fractionation protocol as previously described (Hallett et al., 2008; Warmus et al., 2014). Sub-dissected tissue samples were bathed and homogenized for 30 seconds in TEVP buffer (10 mM Tris base, 5 mM NaF, 1 mM Na_3_VO_4_, 1 mM EDTA, 1 mM EDTA) with 320 mM sucrose and protease and phosphatase inhibitors. A small volume was saved as whole homogenate (WH). Remaining sample was centrifuged at 800 × g for 10 min at 4°C. The supernatant (S1) was removed, and the pellet (P1) was stored in TEVP + inhibitors. S1 was centrifuged at 9200 × g for 10 minutes at 4°C. The supernatant (S2) was removed and stored. The pellet (P2) was resuspended in TEVP + 32 mM sucrose + inhibitors and centrifuged at 25000 × g for 20 minutes at 4°C. The supernatant (LS1) was removed and stored. The pellet (synaptosome fraction) was resuspended in TEVP+ inhibitors and stored. Protein concentration was determined for all samples by bicinchoninic acid method (Pierce), and Western blots were performed to quantify protein content according to our previously described methods (Herskowitz et al., 2011).

#### Western blotting

Twenty-five micrograms of protein per sample were loaded into each well. Primary antibodies were incubated overnight at 4°C. All primary antibodies were from Cell Signaling Technology and used at 1:250, unless otherwise indicated. Primary antibodies include: ROCK1 (catalog # 45171, Abcam), ROCK2 (catalog #5561, Santa Cruz), LIMK (catalog #3842S), pLIMK (catalog #3841), PAK1 (catalog #2602), pMLC (catalog #3671), and tubulin (Iowa Hybridoma Bank, catalog #AA4.3, 1:500). Tubulin was used as a loading control. Odyssey Image Station (Li-Cor) captured images, and Odyssey Application Software (3.0, Li-Cor) quantified band intensities.

### Experimental design and statistical analysis

The effect of ROCK1 or ROCK2 heterozygosity was determined by comparing mouse behavior, neuronal structure, or biochemistry with ROCK1 or ROCK2 homozygotes, respectively. Experimental power and sample sizes per condition were determined by referencing similar studies (Anderson et al., 2014; Radley et al., 2015). Pair-wise comparisons were separately performed for ROCK1^+/+^ and ROCK1^+/−^ and for ROCK2^+/+^ and ROCK2^+/−^. All analyses were conducted with Prism 6.0 (GraphPad Software, La Jolla, CA). Significance level was set at P < 0.05, and data were reported as mean ± standard error of the mean. For spine density, behavioral or biochemical comparisons, animal means were averaged to generate a genotype mean. Comparison of genotype means was conducted by two-tailed student’s t-test, unless non-normally distributed or otherwise indicated. Comparisons of spine densities and morphology distributions were performed as previously described (Boros et al., 2017). Spines of apical and basal dendrites were analyzed separately. Two-way analysis of variance (ANOVA) was used to compare densities of morphological classes between genotypes (factors: spine class and genotype). Morphological spine characteristics were compared using cumulative frequency distributions of individual spines from each genotype. Normality of distribution was assessed by D’Agostino & Pearson omnibus normality test. If non-normally distributed, a non-parametric two-sample Kolmogorov-Smirnov test compared whether genotypes segregated in each spine morphology distribution. Sholl analysis was performed in Neurolucida Explorer by drawing concentric shells at 10 μm intervals from the soma. Apical and basal arbors were analyzed separately. The arbor lengths and number of intersections for each individual neuron and radius were averaged to generate means for each genotype-radius pair. Two-way ANOVA (factors: genotype and radius from soma) compared arbor length or number of intersections between genotypes.

## Results

### Genetic depletion of ROCK1 or ROCK2 induces anxiety-like behaviors

Adult ROCK1^+/−^ and ROCK2^+/−^ mice displayed no differences in weight, general health, in-cage behavior or food consumption compared to littermates. Notably, ROCK1^+/+^ and ROCK1^+/−^ mice are littermates and represent a genetic linage, while ROCK2^+/+^ and ROCK2^+/−^ mice are littermates and represent a separate genetic lineage. To test cognition, EPM was used to assess potential anxiety-like behaviors (Hogg, 1996). The amount of time spent in the open arms was reduced significantly in male and female adult ROCK1^+/−^ or ROCK2^+/−^ mice compared to ROCK1^+/+^ or ROCK2^+/+^, respectively (P=0.0024, t(30)=3.318 for ROCK1^+/−^ and P= 0.0486, t(26)=2.069 for ROCK2^+/−^) (Fig. 1 *A, E*). Similar locomotor activity was observed among all genotypes in the Open field test and no differences in working memory was observed in Y-maze (Fig. 1 *B, C, D, F, G, H*). These results indicate that genetic reduction of ROCK1 or ROCK2 induces anxiety without grossly affecting general activity or basic working memory components.

**Figure 1.**
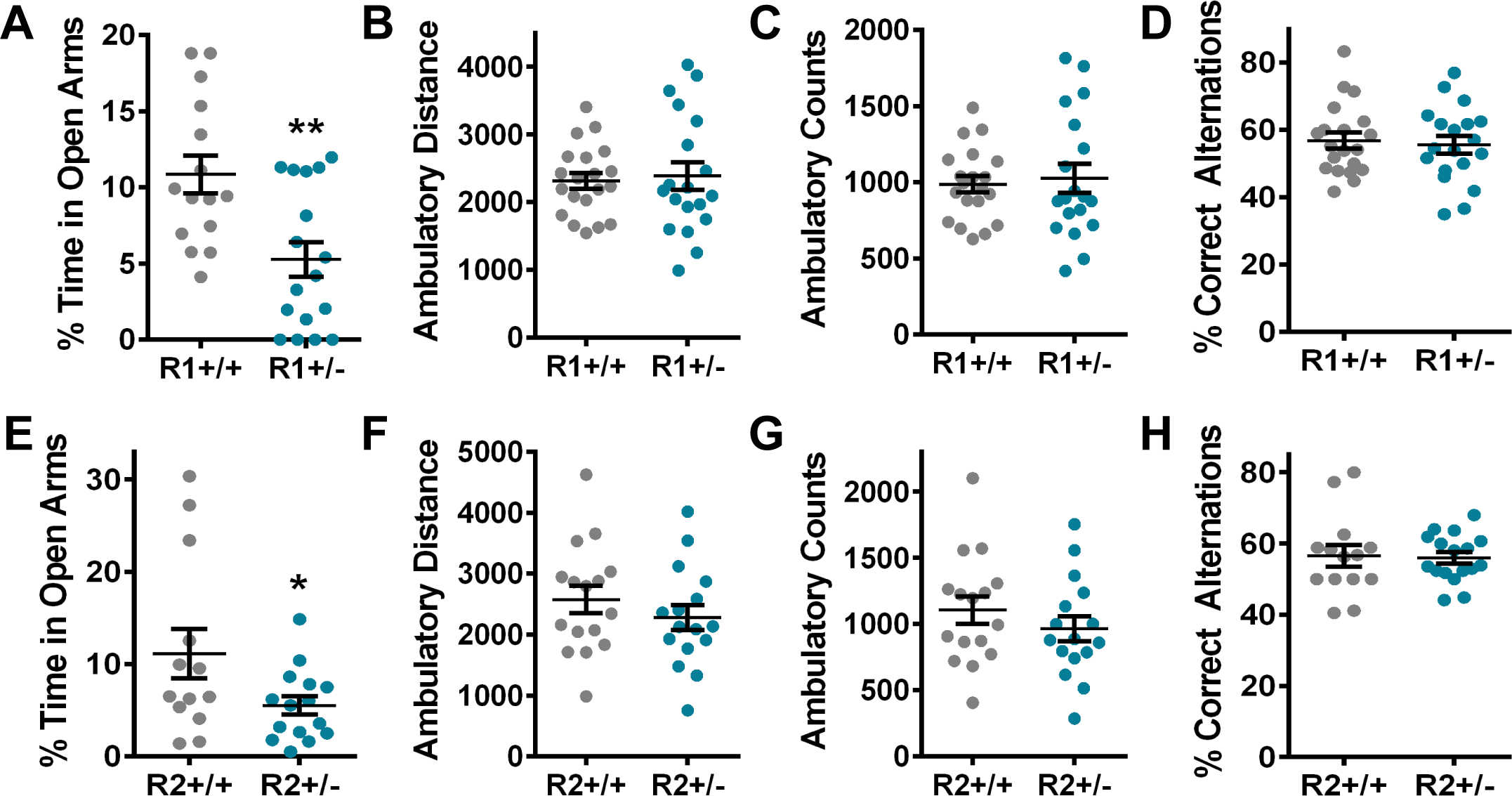
ROCK1 or ROCK2 heterozygosity induces anxiety-like behaviors. All mice were age-matched (4.5-6 months old) and sex-matched. N=15-20 and N=17-19 for ROCK1^+/+^ and ROCK1^+/−^ mice, respectively. N=13-16 and N=15-17 for ROCK2^+/+^ and ROCK2^+/−^ mice, respectively. (A) ROCK1^+/−^ mice spent less time in the open arms of the elevated plus maze (EPM) compared to ROCK1^+/−^ (t-test: **P=0.0024, t=3.318, df=30). (B-C) Open field test for ROCK1^+/+^ and ROCK1^+/−^ mice. (B) Ambulatory distance and (C) ambulatory counts were measured. (D) ROCK1^+/+^ and ROCK1^+/−^ mice performed similarly in Y-maze. (E) ROCK2^+/−^ mice spent less time in the open arms of the EPM compared to ROCK2^+/+^ (t-test: *P=0.0486, t=2.069, df=26). (F-G) Open field test for ROCK2^+/+^ and ROCK2^+/−^ mice show similar (F) ambulatory distance and (G) ambulatory counts. (H) ROCK2^+/+^ and ROCK2^+/−^ mice performed similarly in Y-maze. Lines represent the mean ± standard error of the mean. R1^+/+^, ROCK1^+/+^; R1^+/−,^ ROCK1^+/−^; R2^+/+^, ROCK2^+/+^; R2^+/+^, ROCK2^+/−^.

### ROCK1 reduction alters prefrontal dendritic morphology

The medial prefrontal cortex (mPFC) is required for normal anxiety-related behaviors in the EPM (Shah et al., 2004). Moreover, rodent avoidance of the open arms in the EPM is linked to alterations in the firing rate of single neurons in the mPFC (Adhikari et al., 2011). Based on this, individual layer 2 or 3 pyramidal neurons in the mPFC were targeted for iontophoretic microinjection of the fluorescent dye Lucifer yellow, followed by high-resolution confocal laser scanning microscopy and neuronal 3D reconstructions for morphometry analysis (Fig. 2). A total of 166 dendritic segments from 84 fluorescent dye-filled neurons were imaged for arbor and dendritic spine morphometric analyses (17 ROCK1^+/+^, 19 ROCK1^+/−^, 23 ROCK2^+/+^, 25 ROCK2^+/−^), including 7337, 3845, 4548, 7685 spines in ROCK1^+/+^, ROCK1^+/−^, ROCK2^+/+^, ROCK2^+/−^mice, respectively.

**Figure 2.**
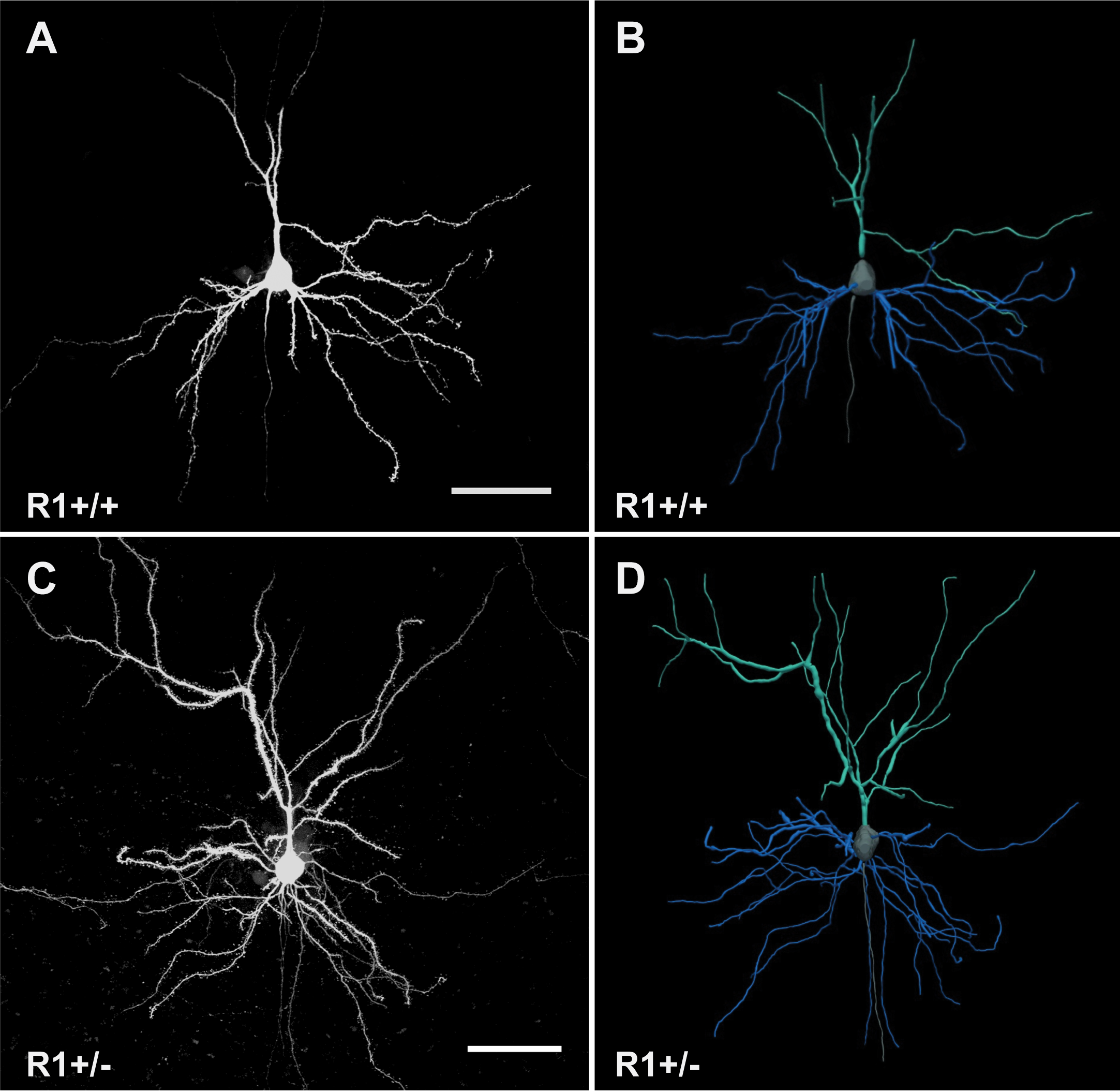
Representative confocal microscope images and corresponding three-dimensional digital reconstruction models of iontophoretically-filled layer 3 pyramidal neurons in the medial prefrontal cortex. Representative maximum-intensity image of (A) ROCK1^+/+^ and (C) ROCK1^+/−^ neurons. Scale bar represents 50 μm. Three-dimensional digital reconstructions of (B) ROCK1^+/+^and (D) ROCK1^+/−^ neurons, corresponding with confocal images. Apical arbors are depicted in teal; basal arbors are shown in blue; and somas are in grey. R1+/+, ROCK1^+/+^; R1+/−, ROCK1^+/−^.

Examination of dendritic length in ROCK1^+/+^ and ROCK1^+/−^ mice by scholl analysis revealed a main effect of genotype by two-way ANOVA on apical and basolateral dendrites (F_(1, 426)_ = 10.89, P = 0.0010 and F_(1,370)_ = 7.083, P = 0.0081, respectively) (Fig. 3 *A, C*). Furthermore, apical and basolateral dendrite intersections were increased in ROCK1^+/−^ mice (F_(1, 426)_ = 10.86, P = 0.0011 and F_(1, 370)_ = 13.08, P = 0.0003, respectively) (Fig. 3 *B, D*). Examination of apical and basolateral dendritic length or intersections in ROCK2^+/+^ and ROCK2^+/−^ mice revealed no statistically significant main effects of genotype (Fig. 3 *E-H*). These data suggest that ROCK1 heterozygosity induces increased dendritic length and complexity at the apical and basal poles of pyramidal neurons in the mPFC. However, neuronal architecture at the dendrite level remains relatively normal with ROCK2 heterozygosity.

**Figure 3.**
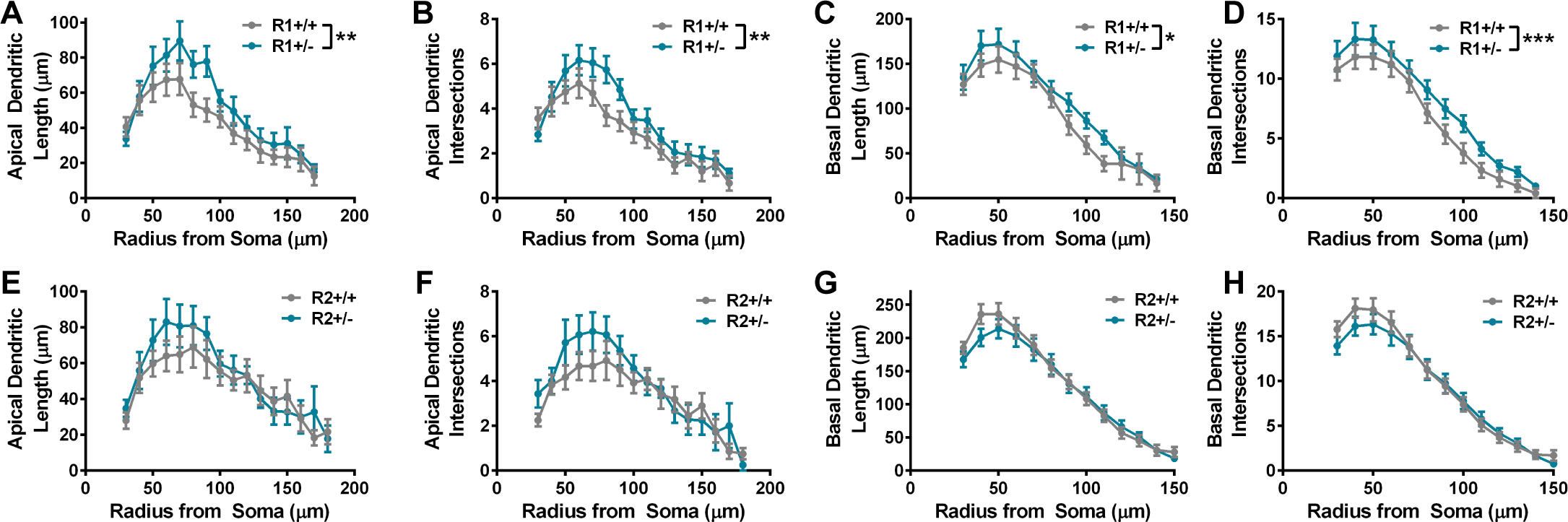
ROCK1 heterozygosity induces increased dendritic length and complexity. Sholl analysis with 10 μm shells was performed for apical and basolateral arbors from (A-D) ROCK1^+/+^ (N=17 neurons from 4 mice) and ROCK1^+/−^ (N=19 neurons from 6 mice) or (E-H) ROCK2^+/+^ (N=23 neurons from 5 mice) and ROCK2^+/−^ (N=25 neurons from 6 mice). Sholl curves for ROCK1^+/+^ and ROCK1^+/−^ indicate that genotypes segregate based on (A) apical dendrite length (Two-way analysis of variance [ANOVA]: **P=0.0010, F_1,426_ = 10.89), (B) apical dendrite intersections (Two-way ANOVA: **P=0.0011, F_1, 426_ = 10.86), (C) basal dendrite length (Two-way ANOVA: **P=0.0081, F_1, 370_ = 7.083), and (D) basal dendrite intersections (Two-way ANOVA: **P = 0.0003, F_1, 370_ = 13.08). Sholl curves for ROCK2^+/+^and ROCK2^+/−^ indicate that genotypes do not segregate based on (E) apical dendrite length, (F) apical dendrite intersections, (G) basal dendrite length, or (H) basal dendrite intersections. Lines represent the mean ± standard error of the mean. R1^+/+^, ROCK1^+/+^; R1^+/−^, ROCK1^+/−^; R2^+/+^, ROCK2^+/+^; R2^+/+^, ROCK2^+/−^.

### Effects of ROCK1 or ROCK2 heterozygosity on prefrontal dendritic spine morphology

*In vitro* and *in vivo* studies indicate that small molecule inhibitors of ROCKs can increase or decrease dendritic spine density, depending on the experimental paradigm, and our previous work demonstrated that pan-ROCK inhibition alters spine morphology in cultured hippocampal neurons (Kang et al., 2009; Hodges et al., 2011; Newell-Litwa et al., 2015; Swanger et al., 2015). However due to the lack of isoform-specificity with pan-ROCK inhibitors, the contribution of ROCK1 or ROCK2 to the observed structural phenotypes could not be determined. Thus, we addressed how genetic depletion of ROCK1 or ROCK2 may affect mPFC dendritic spine morphology. Architectural features of spines were delineated from high-resolution optical stacks of dendrites in fluorescent Lucifer yellow-filled pyramidal neurons using Neurolucida 360 (Fig. 4). Unlike GFP-expressing mice or viral-mediated fluorescent strategies, this technique targets individual neurons and generates information that is comparable to serial section transmission electron microscopy but substantially less prone to the sampling errors common in Golgi-based approaches (Dumitriu et al., 2011).

**Figure 4.**
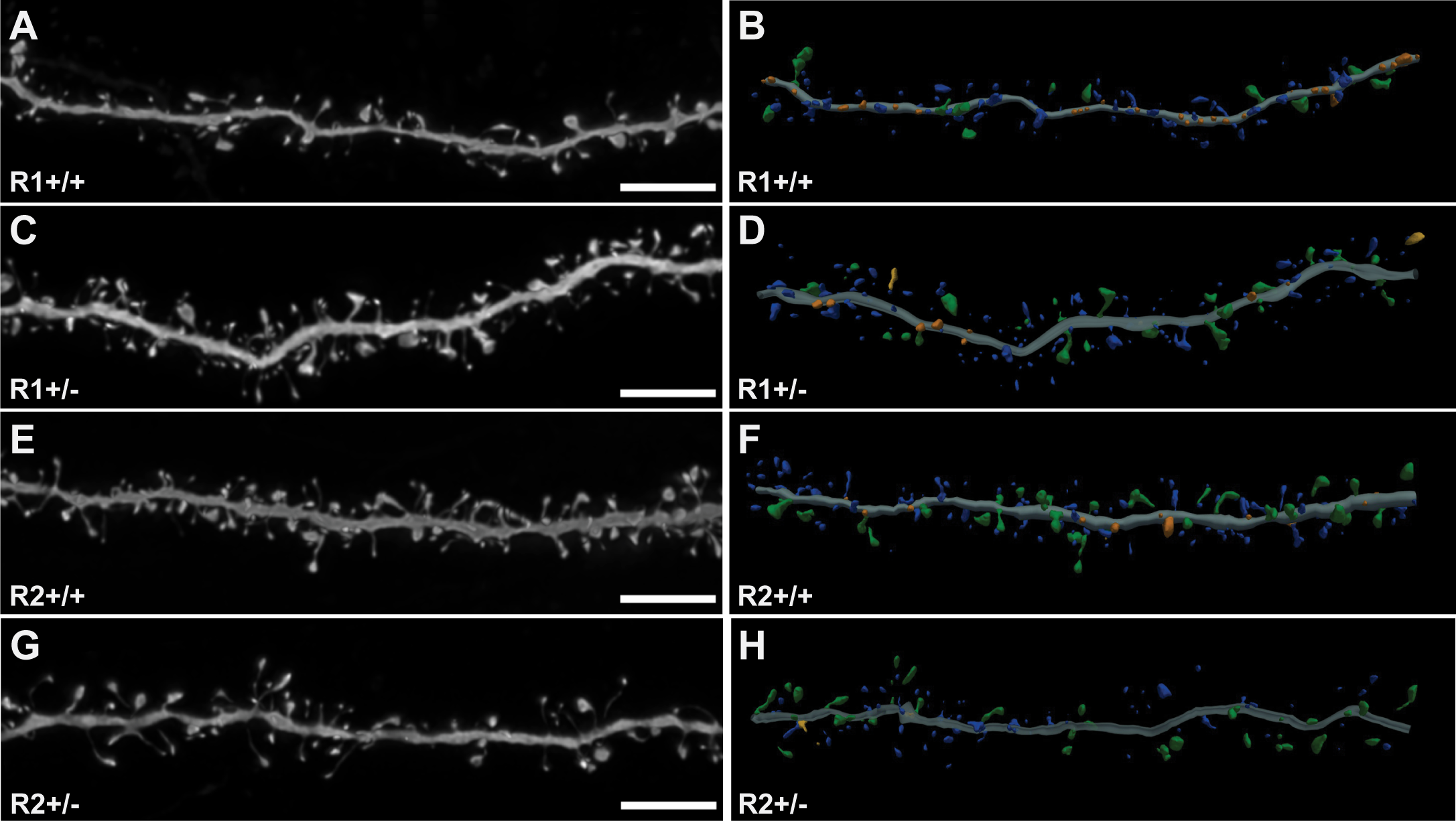
Representative high-resolution confocal microscope images and corresponding three-dimensional digital reconstruction models of dendrites. (A, C, E, G) Representative maximum-intensity confocal images of dye-filled dendrites after deconvolution. Scale bars represent 5 μm. (B, D, F, H) Three-dimensional digital reconstructions of dendrites depicted on the left. Colors correspond to dendritic protrusion classes: blue represents thin spines; orange, stubby spines; green, mushroom spines; and yellow, dendritic filopodia. R1+/+, ROCK1^+/+^; R1+/−, ROCK1^+/−^; R2+/+, ROCK2^+/+^; R2+/+, ROCK2^+/−^.

Spine morphology influences excitatory neurotransmission and synaptic plasticity, and spines can be classified on the basis of their three-dimensional structure as stubby, mushroom, or thin (Harris et al., 1992;Hering and Sheng, 2001; Hayashi and Majewska, 2005). Comparison of dendritic spine density on apical and basolateral dendrites among ROCK1^+/+^ and ROCK1^+/−^ mice revealed no main effect of genotype by two-way ANOVA. This analysis included thin, stubby, and mushroom spine populations (Fig. 5A). To further analyze spine structure, the cumulative distribution of apical and basolateral spine extents (length), head dimeters, and neck diameters for ROCK1^+/+^ and ROCK1^+/−^ mice were plotted. The cumulative frequency plots of apical spines indicated that extent was decreased in ROCK1^+/−^ mice (Kolmogorov-Smirnov: D=0.06705, P<0.0001), whereas head diameter and neck diameter were similar to ROCK1^+/+^ (Fig. 5 *B-D*). The cumulative frequency plots of basal spines indicated that extent was slightly decreased in ROCK1^+/−^ mice (Kolmogorov-Smirnov: D=0.0465, P=0.0372) (Fig. 5E). Head diameter was reduced significantly in ROCK1^+/−^ mice (Kolmogorov-Smirnov: D=0.1382, P<0.0001) (Fig. 5F), and this effect was largely driven by robust decreases in thin spine head diameters (Kolmogorov-Smirnov: D=0.1304, P<0.0001) (Fig. 5G). Neck diameter was also decreased significantly in ROCK1^+/−^ mice (Kolmogorov-Smirnov: D=0.0879, P<0.0001) (Fig. 5H), and this effect was driven by mushroom spine populations (Kolmogorov-Smirnov: D=0.2416, P<0.0001) (Fig. 5I).

**Figure 5.**
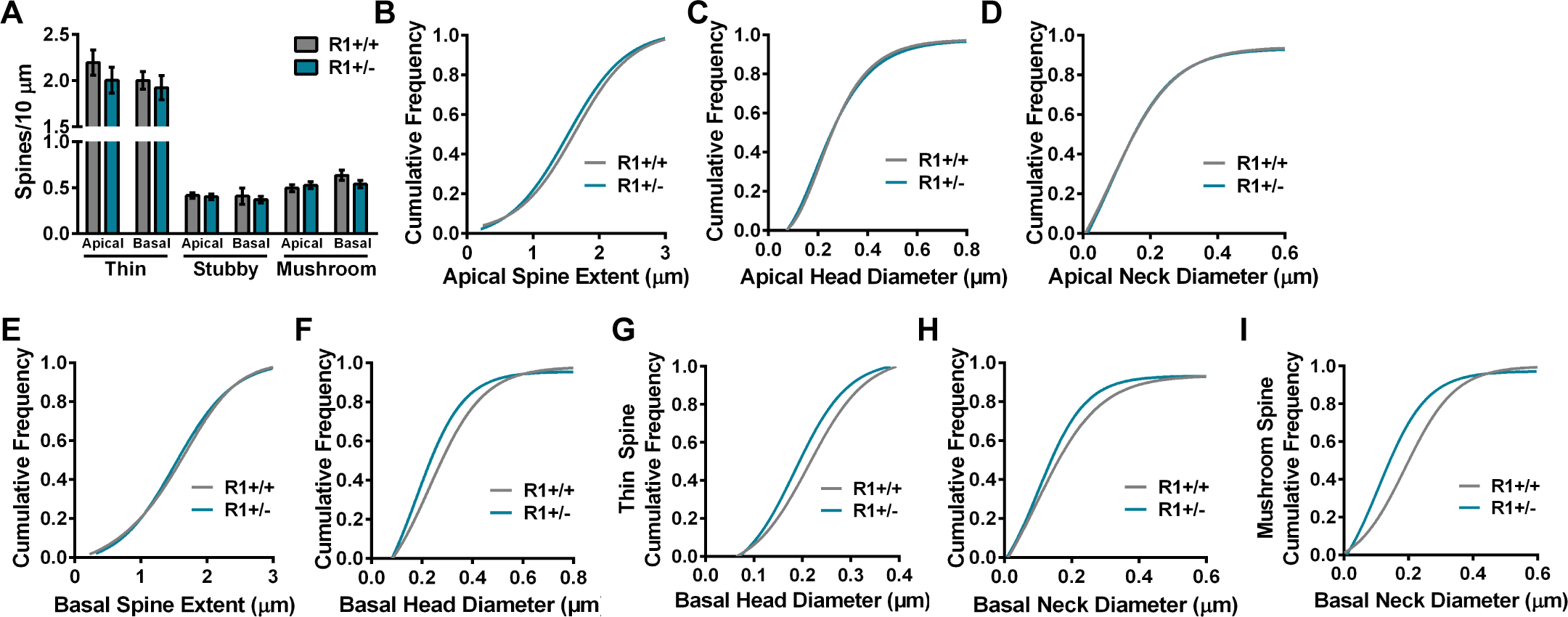
Comparison of apical and basal spine densities and morphology in ROCK1^+/+^ and ROCK1^+/−^ mice. N=34 dendrites and 7337 spines from 5 ROCK1^+/+^ mice and N=38 dendrites and 3845 spines from 6 ROCK1^+/−^ mice. (A) Mean number of thin, stubby, or mushroom spines per 10 μm of apical or basal dendrite was similar between ROCK1^+/+^ and ROCK1^+/−^. (B-I) The cumulative frequency distributions for individual spines were plotted for each genotype and morphological parameter. Comparison of apical spine distribution plots indicate that ROCK1^+/+^ and ROCK1^+/−^ segregate based on (B) apical spine extent (Kolmogorov-Smirnov [KS]: D=0.0671, P<0.0001); but not (C) apical head diameter or (D) apical neck diameter. Comparison of basal spine distribution plots show that genotypes segregate based on (E) basal spine extent (KS: D=0.0465, P=0.0372); (F) basal head diameter (KS: D=0.1382, P<0.0001); (G) basal thin spine head diameter (KS: D=0.1304, P<0.0001); (H) basal neck diameter (KS: D=0.0879, P<0.0001); and (I) basal mushroom spine neck diameter (KS: D=0.2416, P<0.0001). Lines represent the mean ± standard error of the mean. R1^+/+^, ROCK1^+/+^; R1^+/−^, ROCK1^+/−^; R2^+/+^, ROCK2^+/+^; R2+/+, ROCK2^+/−^.

Examination of dendritic spine density, including thin, stubby, and mushroom spine populations, on apical and basolateral dendrites among ROCK2^+/+^ and ROCK2^+/−^ mice revealed no main effect of genotype by two-way ANOVA (Fig. 6A). To analyze spine structure, the cumulative distribution of apical and basolateral spine extents, head dimeters, and neck diameters were plotted. The cumulative frequency plots of apical spines indicated that extent was increased in ROCK2^+/−^ mice (Kolmogorov-Smirnov: D=0.0702, P<0.0001), predominantly driven by increased extent of mushroom spines (Kolmogorov-Smirnov: D=0.1195, P=0.0063) (Fig. 6 *B-C*). Apical spine head and neck diameter were reduced significantly in ROCK2^+/−^ mice (Kolmogorov-Smirnov: D=0.0610, P=0.0002; and Kolmogorov-Smirnov: D=0.0778, P<0.0001, respectively) (Fig. 6 *D-E*). Thin, stubby, and mushroom spine populations contributed approximately equivalently to the effects on head and neck diameter in ROCK2^+/−^ mice. The cumulative frequency plots of basal spines indicated that extent was similar in ROCK2^+/+^ and ROCK2^+/−^ mice, whereas head and neck diameter were reduced in ROCK2^+/−^ mice (Kolmogorov-Smirnov: D=0.0567, P<0.0001; and Kolmogorov-Smirnov: D=0.0541, P=0.0002, respectively) (Fig. 6 *F-H*). Notably, thin spines drove the reduction in neck diameter (Kolmogorov-Smirnov: D=0.0653, P=0.0003), while all spine classes contributed to decreased head diameter in ROCK2^+/−^ mice (Fig. 6I).

**Figure 6.**
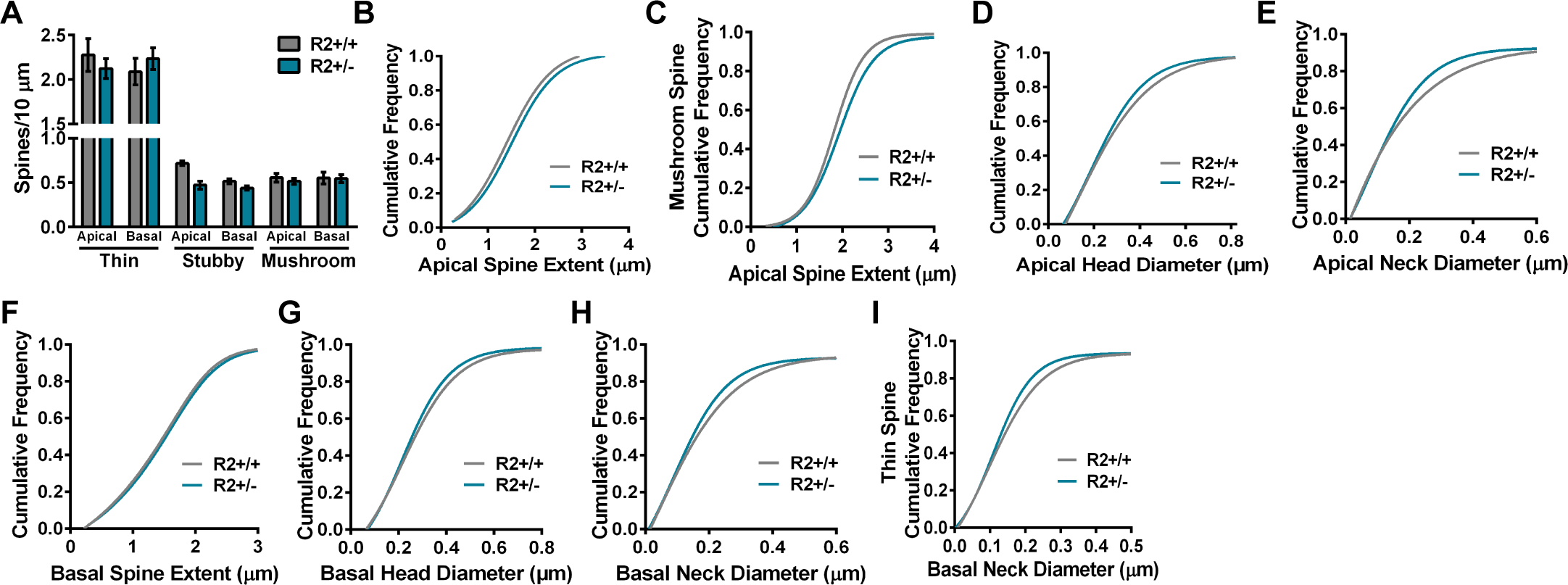
Comparison of apical and basal spine densities and morphology in ROCK2^+/+^ and ROCK2^+/−^. N=46 dendrites and 4548 spines from 4 ROCK2^+/+^ mice and N=50 dendrites and 7685 spines from 5 ROCK2^+/−^ mice.(A) Mean number of thin, stubby, or mushroom spines per 10 μm of apical or basal dendrite was similar between ROCK2^+/+^ and ROCK2^+/−^. (B-I) The cumulative frequency distributions for individual spines were plotted for each genotype and morphological parameter. Comparison of apical spine distribution plots indicate that ROCK2^+/+^ and ROCK2^+/−^ segregate based on (B) apical spine extent (Kolmogorov-Smirnov [KS]: D=0.0702, P<0.0001); (C) apical mushroom spine extent (KS: D=0.1195, P=0.0063); (D) apical head diameter (KS: D=0.0610, P=0.0002); and (E) apical neck diameter (KS: D=0.0778, P<0.0001). (F-I) Comparison of basal spine distribution plots show that genotypes segregate based on (G) basal head diameter (KS: D=0.0567, P<0.0001); (H) basal neck diameter (KS: D=0.0541, P=0.0002); and (I) basal thin spine neck diameter (KS: D=0.0653, P=0.0003); but not (F) basal spine extent. Lines represent the mean ± standard error of the mean. R1^+/+^, ROCK1^+/+^; R1^+/−^, ROCK1^+/−^; R2^+/+^, ROCK2^+/+^; R2^+/+^, ROCK2^+/−^.

Collectively, these findings reveal that apical spine extent is decreased in ROCK1^+/−^ but increased in ROCK2^+/−^ mice. Yet similar reductions in head and neck diameter were observed in apical and basolateral spine populations from ROCK1^+/−^ or ROCK2^+/−^ mice compared to ROCK1^+/+^ or ROCK2^+/+^, respectively. These data support the hypothesis that both isoforms of ROCK play a role in spine head and neck diameter maintenance, whereas ROCK1 and ROCK2 form a yin and yang to regulate spine extent.

### Biochemical analyses of ROCK substrates in synaptic fractions from ROCK1^+/−^ and ROCK2^+/−^ mice

Because ROCK1^+/−^ and ROCK2^+/−^ mice displayed substantial alterations in mPFC dendritic spine morphology, we explored whether there were biochemical changes in key ROCK substrates at synapses. Expectedly, levels of ROCK1 or ROCK2 were decreased approximately 50% in whole homogenates (WH) and synaptosome fractions in heterozygous mice compared to ROCK1^+/+^ and ROCK2^+/+^ (Fig. 7). Previous reports strongly suggest that ROCKs can regulate actin filaments in dendritic spines by phosphorylating LIMKs at threonine 505 and/or threonine 508 (pLIMK) (Sumi et al., 2001; Bosch et al., 2014). Levels of pLIMK were decreased significantly in synaptosome fractions in ROCK1^+/−^ and ROCK2^+/−^ mice compared to ROCK1^+/+^ and ROCK2^+/+^, respectively (P=0.0265, t(10)=2.600 for ROCK1^+/−^ and P=0.0253, t(12)=2.554 for ROCK2^+/−^) (Fig. 7 *B, C, E, F*).In contrast, pLIMK levels were increased significantly in WH in ROCK1^+/−^ and ROCK2^+/−^ mice compared to ROCK1^+/+^ and ROCK2^+/+^, respectively (P=0.0355, t(10)=2.429 for ROCK1^+/−^ and P=0.0004, t(11)=5.000 for ROCK2^+/−^) (Fig. 7 *A, C, D, F*). LIMK protein levels were similar among all genotypes in WH and synaptic fractions, indicating changes in pLIMK were not the result of LIMK fluctuations (Fig. 7 *A, B, D, E*). ROCKs can regulate actin-myosin contractility by phosphorylating MLC at serine 19 (pMLC), and MLC can play a critical role in dendritic spine morphology and synaptic function (Amano et al., 1996; Ryu et al., 2006). Levels of pMLC were decreased significantly in synaptosome fractions of ROCK1^+/−^ mice compared to ROCK1^+/+^ (P=0.0130, t(9)=3.089 for ROCK1^+/−^) but similar in ROCK2^+/+^ and ROCK2^+/−^ mice (Fig. 7 *B, C, E, F*). These results indicate that heterozygosity of either ROCK isoform decreases pLIMK in synapses, correlating to reductions in dendritic spine head and neck diameter in the mPFC (Figs. 5-6). Notably, changes in pMLC were selective to ROCK1^+/−^ mice, suggesting a putative mechanism to explain spine extent reduction in these animals.

**Figure 7.**
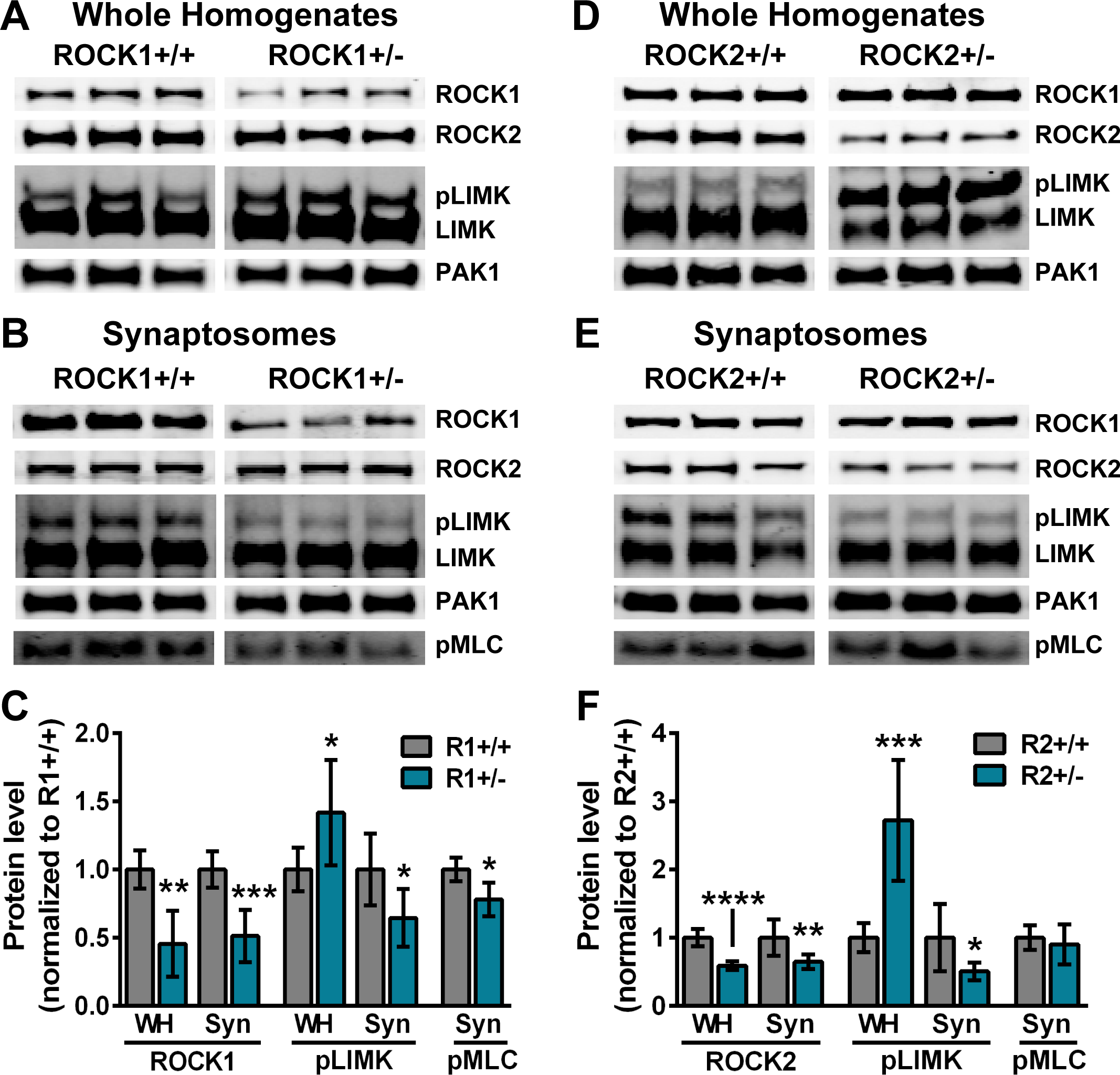
ROCK1 or ROCK2 heterozygosity reduces pLIMK in synaptosome fractions. N=6 mice for all genotypes. Representative Western blots from whole homogenate (WH) and synaptosome (Syn) fractions of (A-B) ROCK1^+/+^ and ROCK1^+/−^ or (D-E) ROCK2^+/+^ and ROCK2^+/−^ mice. For densitometry analysis protein levels of WH or Syn homogenates from ROCK1^+/−^ or ROCK2^+/−^ were normalized to ROCK1^+/+^ or ROCK2^+/+^, respectively. WH and Syn protein levels measured by densitometry were averaged to generate mean per genotype. (C) ROCK1^+/−^ WH showed decreased ROCK1 (t-test: **P=0.0024, t=4.369, df=8) and elevated pLIMK levels (t-test: *P=0.0355, t=2.429, df=10). ROCK1^+/−^ Syn fractions showed decreased ROCK1 (t-test:***P=0.0003, t=5.099, df=11), decreased pLIMK (t-test: *P=0.0265, t=2.600, df=10), and decreased pMLC (t-test: *P=0.0130, t=3.089, df=9). (F) ROCK2^+/−^ WH showed decreased ROCK2 (t-test: ****P<0.0001, t=7.273, df=11) and elevated pLIMK (t-test: ***P=0.0004, t=5.000, df=11). ROCK2^+/−^ Syn fractions showed decreased ROCK2 (t-test: **P=0.0068, t=3.261, df=12) and decreased pLIMK (t-test: *P=0.0253, t=2.554, df=12). Lines represent the mean ± standard error of the mean. WH, whole homogenate; Syn, synaptosome fractions; R1+/+, ROCK1^+/+^; R1+/−, ROCK1^+/−^; R2+/+, ROCK2^+/+^; R2+/+, ROCK2^+/−^.

## Discussion

ROCKs are attractive drug targets, and out of the 29 protein kinase inhibitors that have been used to treat human diseases two are pan-ROCK inhibitors: Fasudil and Ripasudil (Feng et al., 2015). Pharmacologic inhibitor studies have provided much of what we know about ROCKs in the brain, and despite the enormous potential these compounds exhibit to modify human disease-progression, target-selectivity caveats have fogged our view of several fundamental, basic questions. Current ROCK-based therapeutics is moving rapidly toward the development of isoform-specific inhibitors, yet this promising avenue is hampered due to ambiguity over ROCK1- or ROCK2-specific functions in the brain (Julian and Olson, 2014; Feng et al., 2015). Moreover, previous genetic attempts to explore ROCK1 and ROCK2 function were severely limited due to complications of knockout mice homozygosity on mixed genetic backgrounds. To address this, we generated new ROCK1^+/−^ and ROCK2^+/−^ mice on the C57BL/6N background. Through a combination of cognitive behavior, highly optimized three-dimensional modeling of dendritic structure, and biochemistry, our study revealed distinct and complementary roles for ROCK1 and ROCK2 in mPFC structural plasticity.

Although locomotor activity and working memory were normal in mice heterozygous for ROCK1 or ROCK2, both genotypes exhibited anxiety-like behaviors in the EPM compared to ROCK1^+/+^ and ROCK2^+/+^ littermates (Fig. 1). Notably, mice exposed to the pan-ROCK inhibitor Y-27632 did not display anxiety in the EPM or changes in novel object recognition tests but did exhibit enhanced spatial discrimination in Y-maze (Christie et al., 2013). The contrasting outcomes of these pharmacologic and genetic approaches may be attributed to simultaneous inhibition of ROCK1 and ROCK2 or haploinsufficiency of ROCK1 or ROCK2 since birth, respectively. To our knowledge, these are the first reported cognitive examinations that assess the impact of ROCK1 or ROCK2 genetic depletion in rodents. Based on the similarity of EPM phenotypes, these results suggested potential overlap in ROCK1 and ROCK2 function with respect to cognition at the organismal level. However, at the cellular level ROCK1, but not ROCK2, heterozygosity increased dendritic length and complexity of layer 2/3 pyramidal neurons in the mPFC. This implicates ROCK1 as a novel regulatory factor of neuronal dendritic structure. Mirroring our findings in ROCK1 heterozygotes, intracerebroventricular infusion of the pan-ROCK inhibitor Fasudil increased dendrite length in CA1 hippocampal neurons (Couch et al., 2010). Collectively, this suggests Fasudil inhibition of ROCK1 contributed predominantly to the arbor effects; however other studies using Golgi-impregnation and Scholl analysis indicated that homozygous deletion of ROCK2 on the CD-1 background increased dendrite length and intersections in hippocampal neurons (Duffy et al., 2009). We observed no statistically significant changes in dendritic arbors of ROCK2 heterozygous mice, suggesting that more robust depletion of ROCK2 may be needed or that ROCK1 can compensate under experimental conditions of ROCK2 heterozygosity.

Past studies in cultured hippocampal neurons provided a starting point for assessing ROCK1 and ROCK2 specific functions. RNAi depletion of ROCK1 increased spine extent and decreased spine head diameter, whereas ROCK2 knockdown increased head diameter (Newell-Litwa et al., 2015). The findings herein indicated that ROCK1 heterozygosity reduced spine extent but similarly decreased head diameter, while ROCK2 heterozygosity decreased spine head diameter. Moreover, our previous *in vitro* studies revealed that treatment with Y-27632 increased spine density as well as spine extent and width in hippocampal neurons (Swanger et al., 2015). These contrasting effects could represent *in vitro* versus *in vivo* conditions or differences in cell-type (hippocampal versus mPFC pyramidal neurons). The similar global reductions in head and neck diameter that were observed in mPFC apical and basolateral spine populations from ROCK1^+/−^ and ROCK2^+/−^ mice suggests that these overlapping phenotypes likely contributed to the equivalent behavioral outcomes in the EPM. Moreover, both ROCK1^+/−^ and ROCK2^+/−^ mice displayed decreased pLIMK in synaptosome fractions, which may explain the reductions in spine head and neck diameter. Similarly, older studies demonstrated that genetic deletion of LIMK reduced spine head area in hippocampal neurons (Meng et al., 2002). The observed increase in WH pLIMK is more challenging to explain. It could reflect a dysregulation of alternative Rho-GTPase signaling, such as PAK1-mediated phosphorylation of LIMK (Edwards et al., 1999), under conditions of ROCK1 or ROCK2 heterozygosity, but we did not observe changes in PAK1 among genotypes or biochemical fractions (Fig. 7 *A, B, D, E*). Alternatively, increased WH pLIMK may be explained by compensatory phosphorylation by ROCK1 or ROCK2 when the other isoform is depleted. That ROCK1 and ROCK2 appear as a yin and yang to regulate spine extent is a fascinating observation and correlates with previous cultured neuron experiments and pan-ROCK inhibitors, demonstrating either increased or decreased spine extent (Kang et al., 2009; Newell-Litwa et al., 2015; Swanger et al., 2015) (Figs. 5-6). It is possible that pharmacologic pan-ROCK inhibition could reflect ROCK-isoform specific effects depending on the balance of ROCK1 and ROCK2 expression in the cellular system.

Concomitant alternations in dendritic spine extent and head diameter among apical and basolateral spine populations in ROCK1^+/−^ and ROCK2^+/−^ mice may reflect more rapid plasticity to maintain information storage (Grutzendler et al., 2002). Notably, exposure to Y-27632 increased the magnitude of long-term potentiation in adult rat brain hippocampal slices (O’Kane et al., 2004). ROCK1- and ROCK2-mediated structural plasticity of spine neck diameter is equivalently intriguing given the recent demonstration that neck structural plasticity regulates excitatory postsynaptic potential while controlling the biochemical compartmentalization of the overall spine shape (Tonnesen et al., 2014). Collectively, our findings highlight distinct and complementary roles for ROCK1 and ROCK1 in prefrontal cortex structural plasticity and provide a fundamental basis for future development of isoform-selective ROCK inhibitors to treat neurologic disorders.

## Conflict of interest

The authors declare no competing financial interests.

## Acknowledgements

This work was supported by the National Institutes of Health through NIA AG054719 to J.H.H. and NIA AG043552-05 to J.H.H. Additional support stemmed from a New Investigator Research Grant 2015-NIRG-339422 to J.H.H. from the Alzheimer’s Association.

## References

Adhikari A, Topiwala MA, Gordon JA (2011) Single units in the medial prefrontal cortex with anxiety-related firing patterns are preferentially influenced by ventral hippocampal activity. Neuron 71:898–910.

Albersen M, Shindel AW, Mwamukonda KB, Lue TF (2010) The future is today: emerging drugs for the treatment of erectile dysfunction. Expert Opin Emerg Drugs 15:467–480.

Amano M, Ito M, Kimura K, Fukata Y, Chihara K, Nakano T, Matsuura Y, Kaibuchi K (1996) Phosphorylation and activation of myosin by Rho-associated kinase (Rho-kinase). J Biol Chem 271:20246–20249.

Anderson RM, Birnie AK, Koblesky NK, Romig-Martin SA, Radley JJ (2014) Adrenocortical status predicts the degree of age-related deficits in prefrontal structural plasticity and working memory. J Neurosci 34:8387–8397.

Boros BD, Greathouse KM, Gentry EG, Curtis KA, Birchall EL, Gearing M, Herskowitz JH (2017) Dendritic spines provide cognitive resilience against Alzheimer’s disease. Ann Neurol 82:602–614.

Bosch M, Castro J, Saneyoshi T, Matsuno H, Sur M, Hayashi Y (2014) Structural and molecular remodeling of dendritic spine substructures during long-term potentiation. Neuron 82:444–459.

Bradley A et al. (2012) The mammalian gene function resource: the International Knockout Mouse Consortium. Mamm Genome 23:580–586.

Challa P, Arnold JJ (2014) Rho-kinase inhibitors offer a new approach in the treatment of glaucoma. Expert Opin Investig Drugs 23:81–95.

Christie KJ, Turbic A, Turnley AM (2013) Adult hippocampal neurogenesis, Rho kinase inhibition and enhancement of neuronal survival. Neuroscience 247:75–83.

Couch BA, DeMarco GJ, Gourley SL, Koleske AJ (2010) Increased dendrite branching in AbetaPP/PS1 mice and elongation of dendrite arbors by fasudil administration. J Alzheimers Dis 20:1003–1008.

Davies SP, Reddy H, Caivano M, Cohen P (2000) Specificity and mechanism of action of some commonly used protein kinase inhibitors. Biochem J 351:95–105.

DePoy LM, Zimmermann KS, Marvar PJ, Gourley SL (2017) Induction and Blockade of Adolescent Cocaine-Induced Habits. Biol Psychiatry 81:595–605.

Duffy P, Schmandke A, Sigworth J, Narumiya S, Cafferty WB, Strittmatter SM (2009) Rho-associated kinase II (ROCKII) limits axonal growth after trauma within the adult mouse spinal cord. J Neurosci 29:15266–15276.

Dumitriu D, Rodriguez A, Morrison JH (2011) High-throughput, detailed, cell-specific neuroanatomy of dendritic spines using microinjection and confocal microscopy. Nat Protoc 6:1391–1411.

Edwards DC, Sanders LC, Bokoch GM, Gill GN (1999) Activation of LIM-kinase by Pak1 couples Rac/Cdc42 GTPase signalling to actin cytoskeletal dynamics. Nat Cell Biol 1:253–259.

Feng Y, LoGrasso PV, Defert O, Li R (2015) Rho Kinase (ROCK) Inhibitors and Their Therapeutic Potential. J Med Chem.

Gentry EG, Henderson BW, Arrant AE, Gearing M, Feng Y, Riddle NC, Herskowitz JH (2016) Rho Kinase Inhibition as a Therapeutic for Progressive Supranuclear Palsy and Corticobasal Degeneration. J Neurosci 36:1316–1323.

Grutzendler J, Kasthuri N, Gan WB (2002) Long-term dendritic spine stability in the adult cortex. Nature 420:812–816.

Gunther R, Balck A, Koch JC, Nientiedt T, Sereda M, Bahr M, Lingor P, Tonges L (2017) Rho Kinase Inhibition with Fasudil in the SOD1(G93A) Mouse Model of Amyotrophic Lateral Sclerosis-Symptomatic Treatment Potential after Disease Onset. Front Pharmacol 8:17.

Hallett PJ, Collins TL, Standaert DG, Dunah AW (2008) Biochemical fractionation of brain tissue for studies of receptor distribution and trafficking. Curr Protoc Neurosci Chapter 1:Unit 1 16.

Harris KM, Jensen FE, Tsao B (1992) Three-dimensional structure of dendritic spines and synapses in rat hippocampus (CA1) at postnatal day 15 and adult ages: implications for the maturation of synaptic physiology and long-term potentiation. J Neurosci 12:2685–2705.

Hayashi Y, Majewska AK (2005) Dendritic spine geometry: functional implication and regulation. Neuron 46:529–532.

Henderson BW, Gentry EG, Rush T, Troncoso JC, Thambisetty M, Montine TJ, Herskowitz JH (2016) Rho-associated protein kinase 1 (ROCK1) is increased in Alzheimer’s disease and ROCK1 depletion reduces amyloid-beta levels in brain. J Neurochem 138:525–531.

Hering H, Sheng M (2001) Dendritic spines: structure, dynamics and regulation. Nat Rev Neurosci 2:880–888.

Herskowitz JH, Seyfried NT, Gearing M, Kahn RA, Peng J, Levey AI, Lah JJ (2011) Rho kinase II phosphorylation of the lipoprotein receptor LR11/SORLA alters amyloid-beta production. J Biol Chem 286:6117–6127.

Herskowitz JH, Feng Y, Mattheyses AL, Hales CM, Higginbotham LA, Duong DM, Montine TJ, Troncoso JC, Thambisetty M, Seyfried NT, Levey AI, Lah JJ (2013) Pharmacologic inhibition of ROCK2 suppresses amyloid-beta production in an Alzheimer’s disease mouse model. J Neurosci 33:19086–19098.

Hodges JL, Newell-Litwa K, Asmussen H, Vicente-Manzanares M, Horwitz AR (2011) Myosin IIb activity and phosphorylation status determines dendritic spine and post-synaptic density morphology. PLoS One 6:e24149.

Hogg S (1996) A review of the validity and variability of the elevated plus-maze as an animal model of anxiety. Pharmacol Biochem Behav 54:21–30.

Ishizaki T, Maekawa M, Fujisawa K, Okawa K, Iwamatsu A, Fujita A, Watanabe N, Saito Y, Kakizuka A, Morii N, Narumiya S (1996) The small GTP-binding protein Rho binds to and activates a 160 kDa Ser/Thr protein kinase homologous to myotonic dystrophy kinase. EMBO J 15:1885–1893.

Julian L, Olson MF (2014) Rho-associated coiled-coil containing kinases (ROCK): structure, regulation, and functions. Small GTPases 5:e29846.

Kang MG, Guo Y, Huganir RL (2009) AMPA receptor and GEF-H1/Lfc complex regulates dendritic spine development through RhoA signaling cascade. Proc Natl Acad Sci U S A 106:3549–3554.

Kimura K, Ito M, Amano M, Chihara K, Fukata Y, Nakafuku M, Yamamori B, Feng J, Nakano T, Okawa K, Iwamatsu A, Kaibuchi K (1996) Regulation of myosin phosphatase by Rho and Rho-associated kinase (Rho-kinase). Science 273:245–248.

Koch JC, Tonges L, Barski E, Michel U, Bahr M, Lingor P (2014) ROCK2 is a major regulator of axonal degeneration, neuronal death and axonal regeneration in the CNS. Cell Death Dis 5:e1225.

Komers R, Oyama TT, Beard DR, Tikellis C, Xu B, Lotspeich DF, Anderson S (2011) Rho kinase inhibition protects kidneys from diabetic nephropathy without reducing blood pressure. Kidney Int 79:432–442.

Lee DH, Shi J, Jeoung NH, Kim MS, Zabolotny JM, Lee SW, White MF, Wei L, Kim YB (2009) Targeted disruption of ROCK1 causes insulin resistance in vivo. J Biol Chem 284:11776–11780.

Leung T, Manser E, Tan L, Lim L (1995) A novel serine/threonine kinase binding the Ras-related RhoA GTPase which translocates the kinase to peripheral membranes. J Biol Chem 270:29051–29054.

Leung T, Chen XQ, Manser E, Lim L (1996) The p160 RhoA-binding kinase ROK alpha is a member of a kinase family and is involved in the reorganization of the cytoskeleton. Mol Cell Biol 16:5313–5327.

Li Z, Hall AM, Kelinske M, Roberson ED (2014) Seizure resistance without parkinsonism in aged mice after tau reduction. Neurobiol Aging 35:2617–2624.

Matsui T, Amano M, Yamamoto T, Chihara K, Nakafuku M, Ito M, Nakano T, Okawa K, Iwamatsu A, Kaibuchi K (1996) Rho-associated kinase, a novel serine/threonine kinase, as a putative target for small GTP binding protein Rho. EMBO J 15:2208–2216.

Meng Y, Zhang Y, Tregoubov V, Janus C, Cruz L, Jackson M, Lu WY, MacDonald JF, Wang JY, Falls DL, Jia Z (2002) Abnormal spine morphology and enhanced LTP in LIMK-1 knockout mice. Neuron 35:121–133.

Nakagawa O, Fujisawa K, Ishizaki T, Saito Y, Nakao K, Narumiya S (1996) ROCK-I and ROCK-II, two isoforms of Rho-associated coiled-coil forming protein serine/threonine kinase in mice. FEBS Lett 392:189–193.

Newell-Litwa KA, Badoual M, Asmussen H, Patel H, Whitmore L, Horwitz AR (2015) ROCK1 and 2 differentially regulate actomyosin organization to drive cell and synaptic polarity. J Cell Biol 210:225–242.

O’Kane EM, Stone TW, Morris BJ (2004) Increased long-term potentiation in the CA1 region of rat hippocampus via modulation of GTPase signalling or inhibition of Rho kinase. Neuropharmacology 46:879–887.

Olson MF (2008) Applications for ROCK kinase inhibition. Curr Opin Cell Biol 20:242–248.

Radley JJ, Anderson RM, Cosme CV, Glanz RM, Miller MC, Romig-Martin SA, LaLumiere RT (2015) The Contingency of Cocaine Administration Accounts for Structural and Functional Medial Prefrontal Deficits and Increased Adrenocortical Activation. J Neurosci 35:11897–11910.

Rath N, Olson MF (2012) Rho-associated kinases in tumorigenesis: re-considering ROCK inhibition for cancer therapy. EMBO Rep 13:900–908.

Ryu J, Liu L, Wong TP, Wu DC, Burette A, Weinberg R, Wang YT, Sheng M (2006) A critical role for myosin IIb in dendritic spine morphology and synaptic function. Neuron 49:175–182.

Schaafsma D, Gosens R, Zaagsma J, Halayko AJ, Meurs H (2008) Rho kinase inhibitors: a novel therapeutical intervention in asthma? Eur J Pharmacol 585:398–406.

Shah AA, Sjovold T, Treit D (2004) Inactivation of the medial prefrontal cortex with the GABAA receptor agonist muscimol increases open-arm activity in the elevated plus-maze and attenuates shock-probe burying in rats. Brain Res 1028:112–115.

Shi J, Wu X, Surma M, Vemula S, Zhang L, Yang Y, Kapur R, Wei L (2013) Distinct roles for ROCK1 and ROCK2 in the regulation of cell detachment. Cell Death Dis 4:e483.

Shibuya M, Hirai S, Seto M, Satoh S, Ohtomo E (2005) Effects of fasudil in acute ischemic stroke: results of a prospective placebo-controlled double-blind trial. J Neurol Sci 238:31–39.

Shimizu Y, Thumkeo D, Keel J, Ishizaki T, Oshima H, Oshima M, Noda Y, Matsumura F, Taketo MM, Narumiya S (2005) ROCK-I regulates closure of the eyelids and ventral body wall by inducing assembly of actomyosin bundles. J Cell Biol 168:941–953.

Sumi T, Matsumoto K, Nakamura T (2001) Specific activation of LIM kinase 2 via phosphorylation of threonine 505 by ROCK, a Rho-dependent protein kinase. J Biol Chem 276:670–676.

Swanger SA, Mattheyses AL, Gentry EG, Herskowitz JH (2015) ROCK1 and ROCK2 inhibition alters dendritic spine morphology in hippocampal neurons. Cell Logist 5:e1133266.

Swanson AM, DePoy LM, Gourley SL (2017) Inhibiting Rho kinase promotes goal-directed decision making and blocks habitual responding for cocaine. Nat Commun 8:1861.

Tatenhorst L, Eckermann K, Dambeck V, Fonseca-Ornelas L, Walle H, Lopes da Fonseca T, Koch JC, Becker S, Tonges L, Bahr M, Outeiro TF, Zweckstetter M, Lingor P (2016) Fasudil attenuates aggregation of alpha-synuclein in models of Parkinson’s disease. Acta Neuropathol Commun 4:39.

Thumkeo D, Keel J, Ishizaki T, Hirose M, Nonomura K, Oshima H, Oshima M, Taketo MM, Narumiya S (2003) Targeted disruption of the mouse rho-associated kinase 2 gene results in intrauterine growth retardation and fetal death. Mol Cell Biol 23:5043–5055.

Tonnesen J, Katona G, Rozsa B, Nagerl UV (2014) Spine neck plasticity regulates compartmentalization of synapses. Nat Neurosci 17:678–685.

Warmus BA, Sekar DR, McCutchen E, Schellenberg GD, Roberts RC, McMahon LL, Roberson ED (2014) Tau-mediated NMDA receptor impairment underlies dysfunction of a selectively vulnerable network in a mouse model of frontotemporal dementia. J Neurosci 34:16482–16495.

Zhang YM, Bo J, Taffet GE, Chang J, Shi J, Reddy AK, Michael LH, Schneider MD, Entman ML, Schwartz RJ, Wei L (2006) Targeted deletion of ROCK1 protects the heart against pressure overload by inhibiting reactive fibrosis. FASEB J 20:916–925.

